# Machine learning identifies phenotypic profile alterations of human dopaminergic neurons exposed to bisphenols and perfluoroalkyls

**DOI:** 10.1101/2023.08.23.554260

**Authors:** Andrea Di Credico, Amélie Weiss, Massimo Corsini, Giulia Gaggi, Barbara Ghinassi, Johannes H. Wilbertz, Angela Di Baldassarre

## Abstract

Parkinson’s disease (PD) is the second most common neurodegenerative disease and is characterized by the loss of midbrain dopaminergic neurons. Endocrine disruptors (EDs) are active substances that interfere with hormonal signaling. Among EDs, bisphenols (BPs) and perfluoroalkyls (PFs) are chemicals leached from plastics and other household products, and humans are unavoidably exposed to these xenobiotics. Data from animal studies suggest that ED exposure may play a role in PD, but data about the effect of BPs and PFs on human models of the nervous system are lacking. Previous studies demonstrated that machine learning (ML) applied to microscopy data can classify different cell phenotypes based on image features. In this study, the effect of BPs and PFs at different concentrations within the real-life exposure range (0.01, 0.1, 1, and 2 μM) on the phenotypic profile of human stem cell-derived midbrain dopaminergic neurons (mDANs) was analyzed. Cells exposed for 72 hours to the xenobiotics were stained with neuronal markers and evaluated using high content microscopy yielding 126 different phenotypic features. Two different ML models (XGBoost and LightGBM) were trained to classify ED-treated versus control mDANs. ED-treated mDANs were identified with high accuracy (0.92). Assessment of the phenotypic feature contribution to the classification showed that EDs induced a significant increase of alpha-synuclein (αSyn) and tyrosine hydroxylase (TH) staining intensity within the neurons. Moreover, microtubule-associated protein 2 (MAP2) neurite length and branching were significantly diminished in treated neurons. Our study shows that human mDANs are adversely impacted by exposure to EDs, causing their phenotype to shift and exhibit more characteristics of PD. Importantly, ML-supported high-content imaging can identify concrete but subtle subcellular phenotypic changes that can be easily overlooked by visual inspection alone and that define EDs effects in mDANs, thus enabling further pathological characterization in the future.

## Introduction

Parkinson’s disease (PD) is the second most common neurodegenerative disease, affecting about 3% of the population above 65 years [1]. The pathogenesis of PD involves a combination of environmental and genetic risk factors, which collectively contribute to the development and progression of the disease. The main cellular and molecular hallmarks of PD are represented by the loss of midbrain dopaminergic neurons (mDANs) in the substantia nigra, and intracellular aggregation of alpha-synuclein (αSyn), respectively [2]. Importantly, αSyn aggregates can disrupt normal cellular processes and contribute to the repression of tyrosine hydroxylase (TH), the rate-limiting enzyme in brain catecholamine biosynthesis, decreasing dopamine production [3]. These features lead to the onset of the characteristic motor (e.g., bradykinesia, rigidity, and tremors) and non-motor (e.g., cognitive impairment, psychiatric disturbances, and sleep disorders) symptoms of PD [4].

Endocrine disruptors (EDs) are hormonally active substances present in the environment, including household and industrial products, and can have adverse effects on human health [5]. EDs include bisphenols (BPs), such as bisphenol A (BPA) and S (BPS), and perfluoroalkyls (PFs), such as perfluorooctanesulfonate (PFOS) and perfluorooctanote (PFOA). These chemicals are widely diffuse, since BPs are used to produce polymers and resins for the production of polycarbonate plastics, food packaging, food cans, and thermal receipts [6]. Similarly, PFs are found in different items of common use as cookware and paper food packaging [7]. As a consequence, humans are constantly and unavoidably exposed to these xenobiotics that may threaten human health via different routes such as dermal absorption, inhalation and dietary ingestion [8] and EDs have been detected in human serum, urine, placental tissue, umbilical cord blood and breast milk [9–11]. These findings underscore the ability of EDs to enter and persist within the human body, raising serious concerns about their detrimental effects on human health. Although the molecular mechanism has not been completely clarified, it is generally accepted that BPs act as xenoestrogen, binding and activating the estrogen receptors (ER) α and β, while PFs can interfere with the ER, the androgen and thyroid hormone receptors [12,13]. Numerous studied have associated exposure to these EDs with a range of health concerns, including reproductive disorders, developmental abnormalities, metabolic dysfunction, and an increased risk of several cancers [5,14,15].

Emerging data indicate that BPs and PFs also negatively affect the nervous system [16– 18]. Exposure to these chemicals may deteriorate the dopaminergic system, suggesting a role in PD development [19]. Numerous research works have contributed to explaining this association. For instance, studies conducted in zebrafish and in *Drosophila melanogaster* have demonstrated that BPs significantly alter the dopaminergic system [16,19]. Similarly, BPA-exposed monkey fetuses display reduced levels of dopamine in midbrain dopaminergic neurons [20] and a recent investigation showed that EDs exacerbated phenotypes in a murine PD model [21]. Although these studies suggest that BPs and PFs have the capacity to alter the dopaminergic system thus contributing to the development and progression of PD, current scientific literature lacks information about the involvement of these xenobiotics on human mDANs pathology. Although epidemiological studies show a relationship between EDs exposure and neurodegenerative diseases [22], it is currently unclear which aspects of human mDAN cellular biology can be modified by BPs and PFs, leaving a critical gap in the understanding of the mechanisms underpinning ED-induced neurotoxicity. In addition, most toxicological studies are performed using high concentrations of EDs in the range of hundreds of μM to several mM, that do not mimic a realistic exposure [23,24]. There is thus the need to find possible causal links between EDs and the onset of neurodegenerative diseases. In recent studies, machine learning (ML) classification approaches have been successfully used for cell line stratification and identification of chemical-treated human mDANs and could be exploited for neurotoxicity studies *in vitro* [25–27].

The primary objective of this study was to examine the pathological impact of exposure to BPs and PFs on human stem cell-derived mDANs, a cell type widely used for disease modelling [28–30]. This was achieved by using high-content fluorescence microscopy to analyze the morphological characteristics affected by these EDs. Our hypothesis was that EDs can induce specific morphological modifications in the phenotypic profile of mDANs similar to that observed in PD patients. For this purpose, human mDANs were treated with increasing low-dose concentrations (0.01, 0.1, 1, and 2 μM) of BPA, BPS, PFOS, and PFOA for 72 hours and stained with specific mDANs markers. We then quantified 126 phenotypic features from the high-content imaging dataset and two different ML models (XGBoost and LightGBM) were trained to classify ED-treated versus control mDANs. By means of this image data-based ML classification approach we measured at which concentrations EDs induce overall phenotypic changes and which neuronal phenotypic features are most impacted.

## Material and Methods

### Preparation of medium and plates

On Day 1, Complete Maintenance Media and plates for neuron seeding were prepared. All reagents are listed in **Table S1**, and compositions of solutions and buffers are described in **Table S2**. For coating, Laminin was diluted 1:10 in cold PBS +/+ and added to each well of a previously PDL coated 384-well plate and incubated overnight at 4°C.

### Neuron cultures and Compound treatment

Commercially available cryopreserved 35 days old human induced pluripotent stem cell (hiPSC) derived mDANs were used for this study (**Table S1**). On Day 0, neurons were thawed and, seeded in a 384 well plate in Complete Maintenance Medium at 15,000 cells/well in a final volume of 60 μL per well according to the manufacturers protocol. Plate edge wells were not used. On Day 3, medium change and compound treatment was performed. BPS, BPA, PFOS, and PFOA were dissolved in methanol and a 1.5X solution for all desired concentrations (0.01, 0.1, 1, and 2 μM) was prepared using Complete Maintenance Medium. As a neutral control, the highest methanol concentration of each tested compound was used. To treat the cells, 40 μL of medium per well were aspirated and substituted with an equal volume of 1.5X compound solution. The treated neurons were then incubated at 37°C and 5% CO_2_ for 72 hours.

### Fixation and staining

Neurons were fixed in 4% PFA for 30 minutes and then permeabilized and blocked in a 1X blocking solution (**Table S2**) for 1 hour at room temperature (RT). Cells were washed, labelled with the primary antibodies diluted in primary staining solution (**Table S2**) overnight at 4°C and stained with the appropriate secondary antibodies for 2 hours at RT. Wells were washed with PBS and nuclei counterstained with Hoechst. All the steps were performed using an automated liquid handler.

### Imaging and analysis

Images were acquired using an automated Yokogawa confocal fluorescence microscope. A 40X objective was used to acquire 16 fields/well using Z-stacks consisting of 3 Z-slices separated by 2 μm. Exposure times and laser intensities were adjusted for each of the 4 fluorescent channels separately to obtain an optimal dynamic range of the fluorescent intensities and prevent signal saturation. Images were stored as TIFF files. Image segmentation and phenotypic feature extraction were required for the creation of quantitative phenotypic profiles. Our in-house developed software PhenoLink was used to extract quantitative information from the multichannel images (**Table S1)**. Image segmentation was performed on illumination corrected raw images based on fluorescent channel intensity thresholds empirically determined per plate. Segmented cells were subdivided into four compartments: whole cell, cytoplasm, membrane, and nucleus (**Figure 3A**). The cell compartment was defined as the total MAP2 staining positive area, while cytoplasm was defined by the cell area minus the nuclear area originating from the Hoechst stain. The membrane compartment was defined by a 5-pixel wide window inside of the cell edges. Based on the segmentations, in total 126 quantitative image features were calculated and averaged per well (**Table S3**). Shape features were computed on the boundaries of segmented compartments and include size and shape metrics. Intensity-based features were computed from intensity values in each channel of the images. Texture features quantify the regularity of intensities in images. Microenvironment or context features include counts and spatial relationships within cells, such as the correlation of two channel intensities.

The resulting quantitative data was then used to construct median phenotypic profiles per treatment condition and to compare phenotypic profiles. A Python-based Jupyter notebook is provided to perform data standardization, supervised classification, and plotting (**Table S1). Figure 1** schematically depicts the experimental workflow.

**Figure 1.**
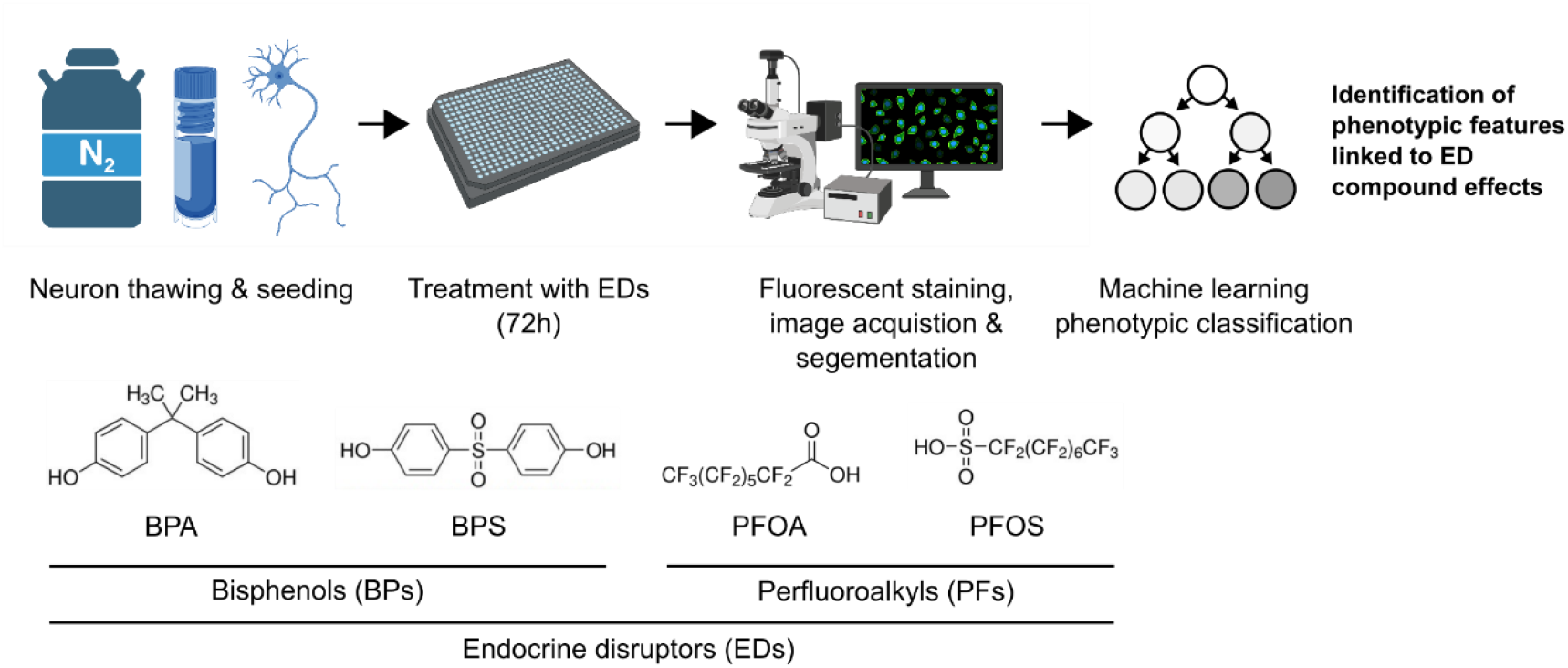
Experimental workflow. Following thawing and seeding, mDANs were treated for 72 hours with the selected EDs at different doses. Then, fluorescent staining against αSyn, TH, MAPs, and DNA was performed. The acquired fluorescent images were segmented and 126 different features were quantified. Finally, ML classifiers were applied to the image-derived data to detect the phenotypic modulations linked to ED effects.

### Data set composition, processing and statistics

Data was generated using two biological replicates representing the separate thawing and culture of mDANs from two cryovials. Within each biological replicate at least three technical replicates (wells) were generated. The used dataset contains 126 columns with continuous phenotypic feature data derived from the image segmentation workflow described above. Additional columns include categorical data that detail the experimental conditions used, such as the position on the plate or the chemical treatment applied. Each row in the dataset represents the mean values per well of a 384-well plate, derived from 16 images. To be suitable for ML, all data was scaled per phenotypic feature using the RobustScaler method in the Python package scikit-learn. RobustScaler scales the data according to the interquartile range (IQR). The IQR is the range between the 1st quartile (25th quantile) and the 3rd quartile (75th quantile). Continuous data was graphically reported by dose-response graphs showing the technical replicate data points, the mean and the 95% confidence interval (CI). Analysis of variance (ANOVA) was used to determine statistically significant differences between the different concentration of each ED on mDAN phenotypical features.

When statistical differences were found, a Tukey post-hoc test was employed. Results were considered significant when p<0.05. We provide a Python-based Jupyter notebook to reproduce all data standardization and plotting steps (**Table S1)**.

### ML classification

In brief, a pipeline was created for preprocessing and classification using the XGBoost or LightGBM classifiers. The data was split into training and testing sets, with 10% of the data being used for testing. Grid search cross-validation with 5 folds was used to find the best hyperparameters for each classifier. The pipeline was then trained on the full training set using the best hyperparameters found. Feature importance were calculated and plotted to show the most important features used by the models. Predictions were made on the test set, and the probabilities of each sample belonging to the “control” class were calculated. The accuracy of the classifier on the test set was evaluated, and a confusion matrix was computed and plotted as a heatmap. 10-fold cross-validation scores were also calculated and plotted across the whole dataset. We provide a Python-based Jupyter notebook to reproduce all data pre-processing, ML, and visualization steps (**Table S1)**.

## Results

### ED treatment increases αSyn staining intensity in human mDANs

mDANs were treated for 72 hours with BPA, BPS, PFOS, and PFOA at four different concentrations (i.e., 0.01, 0.1, 1, and 2 μM) and stained for immunofluorescence analysis. As expected, the imaged neurons were positive for TH and MAP2 and showed the mDAN typical morphology (**Figure 2A**). Methanol only was used as vehicle control (**Figure 2B**). The acquired raw images were segmented, and different phenotypic features were quantified (**Table S3**). Among these, the features “living cells” and “αSyn intensity in TH+ cells” were first considered to evaluate compound toxicity (**Figure 2C**). Living cells were defined as nuclei with a size larger than 2000 pixels and an average pixel intensity lower than 1500. Smaller and brighter nuclei were assumed to show signs of DNA compaction and were considered as apoptotic. No modifications in cell viability were observed when mDANs were treated with BPA, BPS, PFOS or PFOA (**Figure 2D**,), however, a significant difference of αSyn intensity in TH+ cells were found in EDs treated mDANs (**Figure 2D**), except for PFOS (**Figure 2D**). Specifically, αSyn fluorescent intensity was higher in BPA 0.1 μM and 2 μM treated samples, BPS significantly increased αSyn at 2 μM while PFOA was effective in increasing αSyn levels at 0.1, 1, and 2 μM. mDAN imaging therefore indicates that EDs at the tested doses did not affect cell viability but may increase αSyn expression levels.

**Figure 2.**
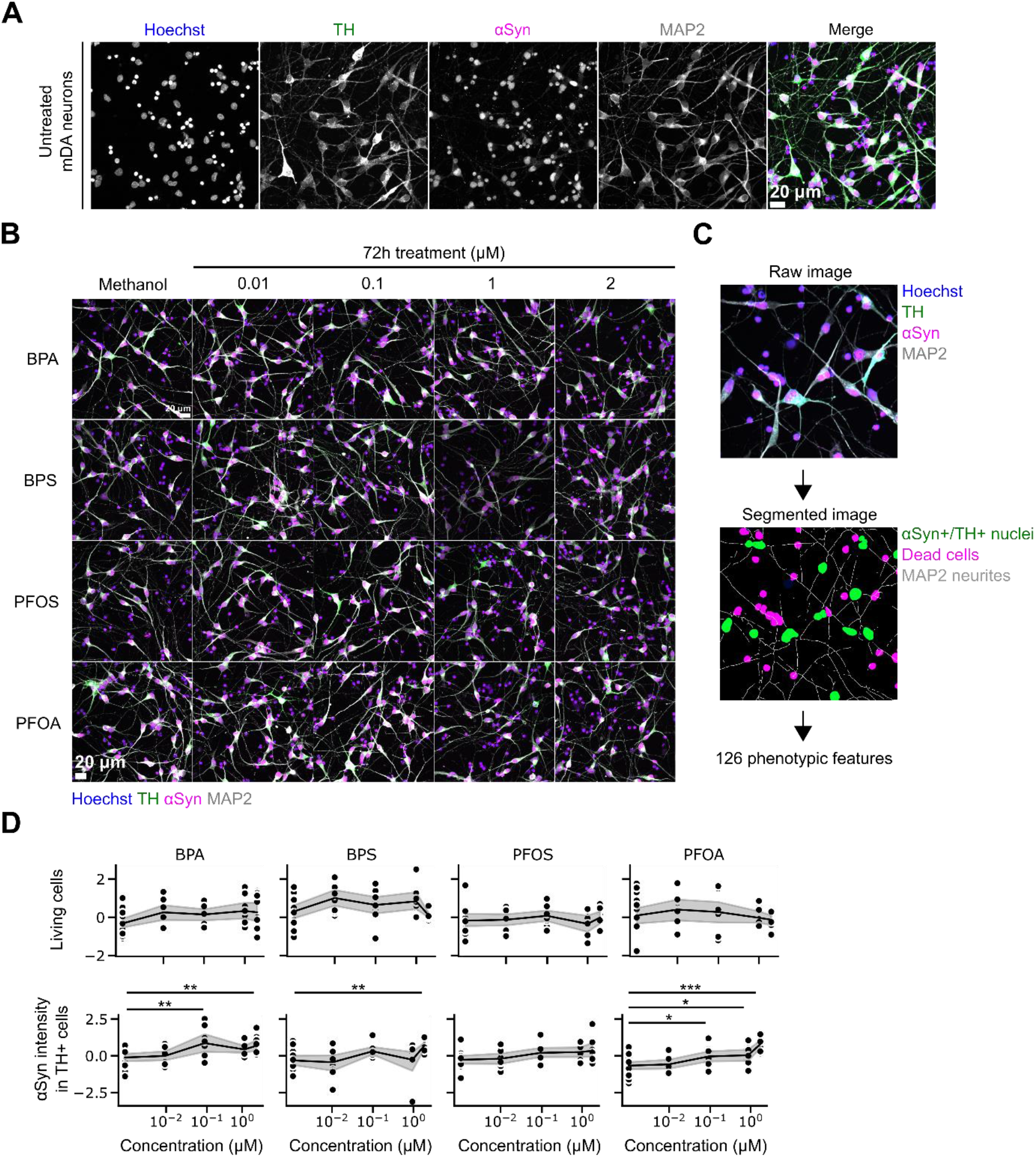
Increased αSyn levels in BP and PF treated human mDANs. (A) After 7 days of culture, mDANs were immunostained against TH, αSyn, and MAP2; nuclei were counterstained with Hoechst. (B) mDANs were treated with increasing concentration (0.01, 0.1, 1, and 2 μM) of four different EDs (BPA, BPS, PFOS, and PFOA) and 16 images were recorded per well. Cells exposed to methanol (vehicle) only were used as control. (C) Example of image segmentation. Raw images from individual channels were segmented, and 126 different phenotypic features describing signal shape, texture, intensity, and localization, were extracted (**Table S3**). (D) Dose-response graphs and statistical analysis describing the effect of treatments on the number of living cells, and αSyn intensity in TH+ neurons. Data is shown as normalized single data points (black dots), mean (black lines) and 95% CI of the mean (gray area). *p<.05, **p<.01, ***p<.001 compared to methanol control.

**Figure 3.**
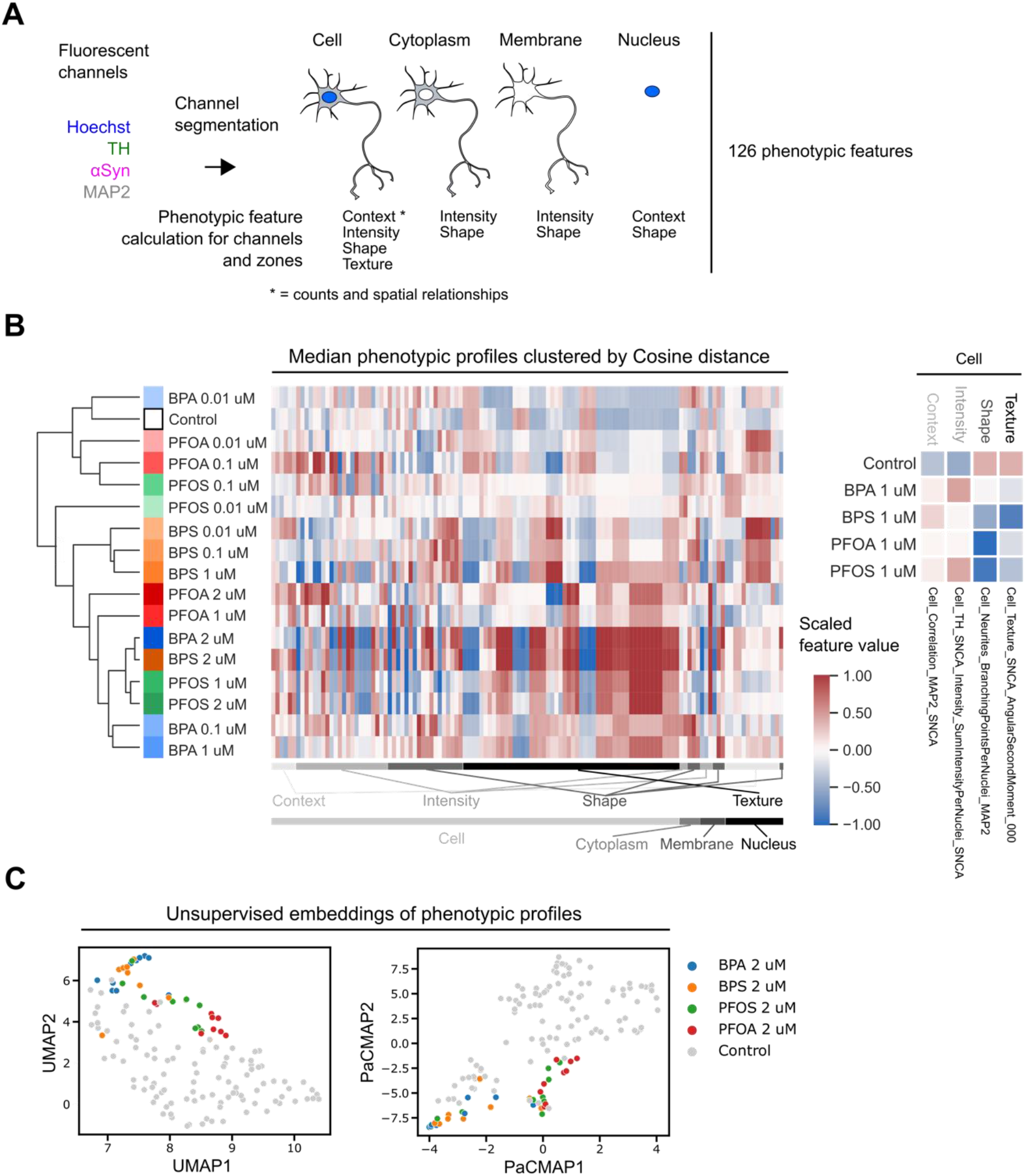
Phenotypic profiles derived from ED-treated mDANs indicate morphological changes in different cellular compartments. (A) From segmented images context, intensity, shape and texture phenotypic features were calculated for different subcellular regions. (B) Phenotypic features were scaled, median aggregated by treatment condition and clustered using the pairwise Cosine distance between each profile. Features were ordered by cellular localization and feature type. Example phenotypic features from each features class illustrate morphological changes after 1 μM ED treatment. (C) UMAP and PaCMAP embeddings with control and 2 μM compound treated per well data.

### BP and PF treatments modify human mDANs phenotypic profile

Humans are usually chronically exposed to low doses of EDs that can induce changes in cellular biology which at least initially may be slight and difficult to detect. We therefore combined 126 morphological features to create a phenotypic profile of mDANs and investigated whether xenobiotic compound exposure would lead to specific signatures. The phenotypic features, originating from the segmented four microscopy channels, were further subdivided into the four compartments: cell, cytoplasm, membrane, and nucleus (**Figure 3A**). Shape, intensity-based and texture features were combined to a treatment-specific phenotypic profile (**Figure 3A, Table S3**). We performed clustering analysis to compare similarities among samples. Each phenotypic profile was considered as a vector and the Cosine distance was measured between profiles. Clustering analysis showed that the xenobiotics, even at low concentrations induced differences in the phenotypic profiles in human mDANs (**Figure 3B**). Lower concentrations tended to cluster closer to the control than higher concentrations indicating that there is a dose-response relationship between increasing ED concentration and variation of the overall phenotypic profiles (**Figure 3B**). Only the 0.01 μM BPS sample was positioned slightly outside the cluster containing other low-concentration EDs and the control. Plotting all profiles also illustrated that many phenotypic features are changed by ED treatment and that not only a single feature class or cellular compartment is affected. Phenotypic profiles also allow to zoom into phenotypic features of interest. Picking one example per features class (context, intensity, shape, texture) clearly showed that 1 μM of all four tested EDs is sufficient to induce observable changes in four selected features describing the correlation between the MAP2 and αSyn channels, the αSyn staining intensity, neurite complexity, and the second angular moment of the αSyn channel texture (**Figure 3B**, right panel).

To further visualize the relationships between the phenotypic profiles, the two embedding techniques Uniform Manifold Approximation and Projection (UMAP) and Pairwise Controlled Manifold Approximation (PaCMAP) were used [28,29] (**Figure 3C**). Embeddings are useful to display high-dimensional data because they can translate high-dimensional data into a relatively low-dimensional space such as a 2D graph. Both embeddings show that phenotypic profiles from ED treated wells group separately from control treated wells (**Figure 3C**). Further, both embeddings show that BPs (BPA and BPS) and PFs (PFOA and PFOS) tend to group together, indicating that the observed morphometric changes are specific to each compound class and related to their chemical structure.

### ML classification discriminates ED-treated mDANs based on image-derived neuronal features

Due to the high dimensionality of phenotypic profiles and the large number of changed features, it is not trivial to identify feature patterns that correlate with an experimental condition. Supervised ML classification algorithms are designed to learn rules from large and complex datasets and compute the probability of a new data point falling into predefined classes, such as ED treated or untreated conditions. We exploited “open” ML classifiers that allowed us to deduce which data features are most explanatory for the observed differences between classes and to identify generalizable rules related to the cell biological effects of ED exposure.

XGBoost and LightGBM are two popular supervised ML algorithms. Due to their use of decision trees, the feature weights of the trained models can be extracted and inform about differences between classes. The main difference between XGBoost and LightGBM is how they build decision trees. XGBoost builds trees one level at a time, while LightGBM focuses on the leaves, or endpoints, of the tree. Briefly, all the image datasets for each condition were divided into a train set (90 % of the dataset) and a test set (10 % of the dataset). XGBoost and LightGBM were then trained to distinguish treated from untreated phenotypic profiles using the training set. All ED concentrations ranging from 0.01 to 2 μM were assigned to the treated class. The trained model was applied to the previously unseen test set phenotypic profile data, in order to cross validate the classification performance (**Figure 4A**).

**Figure 4.**
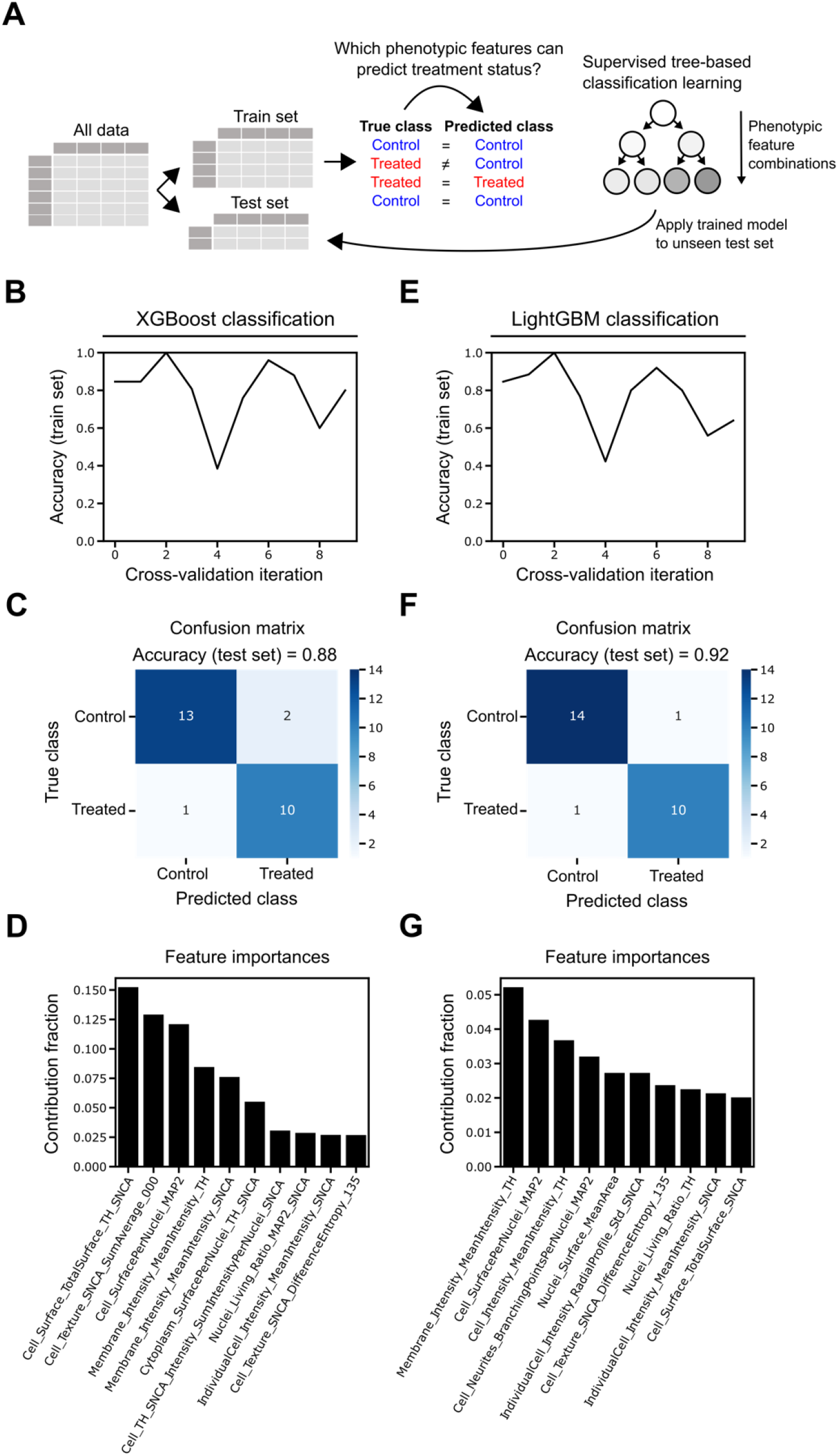
Image data-derived ML classification to predict mDAN phenotypes. (A) Schematic representation of ML training and testing methods used to predict mDAN phenotypes based on treatments. (B, C, D) Classification accuracy for 10 different cross-validation iterations using the training data, confusion matrix graph visually representing the number of times the XGBoost algorithm correctly predicted the experimental condition in the test dataset, and the ten most important features contributing to XGBoost classification performance. (E, F, G) Classification accuracy for 10 different cross-validation iterations using the training data,) confusion matrix graph visually representing the number of times the LightGBM algorithm correctly predicted the experimental condition in the test dataset, and the ten most important features contributing to LightGBM classification performance.

Cross-validation is useful to prevent model overfitting when data is limited and is the application and evaluation of a trained ML model on different slices of the training dataset. When applying 10-fold cross-validation on the training dataset both the XGBoost and LightGBM classification algorithms showed a similar performance in predicting the phenotypic classes (**Figure 4B** and **E**). Due to the relatively small size of the dataset, both XGBoost and LighGBM showed accuracy variations during training from 0.38 to 1.0 (median: 0.83) and 0.4 to 1.0 (median: 0.8), respectively. A confusion matrix was plotted to check for by-class errors and the prediction accuracy of the two different algorithms when applied exclusively to the test set. The test set contained data from 15 control and 11 chemicals treated wells across all concentrations and EDs. XGBoost classification predicted neuronal phenotypes (i.e. methanol control vs. treated) with an accuracy of 0.88 and miss-classified only 2 out 15 control wells as treated and 1 out of 11 treated wells as control wells (**Figure 4C**). LightGBM classification performed similarly and resulted in an accuracy of 0.92 while erroneously classifying only 1 well per treatment category (**Figure 4F**). Next, we extracted the ten most important features involved in the classification performance of the trained models. 6 features contributed between 5-15% to XGBoost model performance, while LightGBM model performance relied on a larger range of features with only a single feature contributing more than 5%. For both classification algorithms, the intensity of TH signal around the cytoplasmic membrane, the MAP2 cell surface per nucleus, and the cellular intensity of αSyn were among the most contributing features to distinguish control and EDs treated samples (**Figure 4D** and **G**).

### Neurite-related features of mDANs are negatively affected by BP and PF treatments

Both classification algorithms indicated that the cellular surface evidenced by the MAP2 signal is a key feature explaining the differences between ED-treated and untreated mDANs (**Figure 4D** and **G**). Upon exposure to the BPs and PFs, mDANs exhibited a visually notable decrease in the length of their neurites (**Figure 5A**) and this negative effect was mostly dose-dependent, being more evident for all EDs at 2 μM (**Figure 5B**, top row). Specifically, Tukey’s post-hoc analysis demonstrated that BPA decreased neurite length at 0.1 and 2 μM. For both, BPS and PFOS, the decrease occurred when mDANs were exposed to 1 and 2 μM. PFOA decreased neurite length at 0.1, 1, and 2 μM.

**Figure 5.**
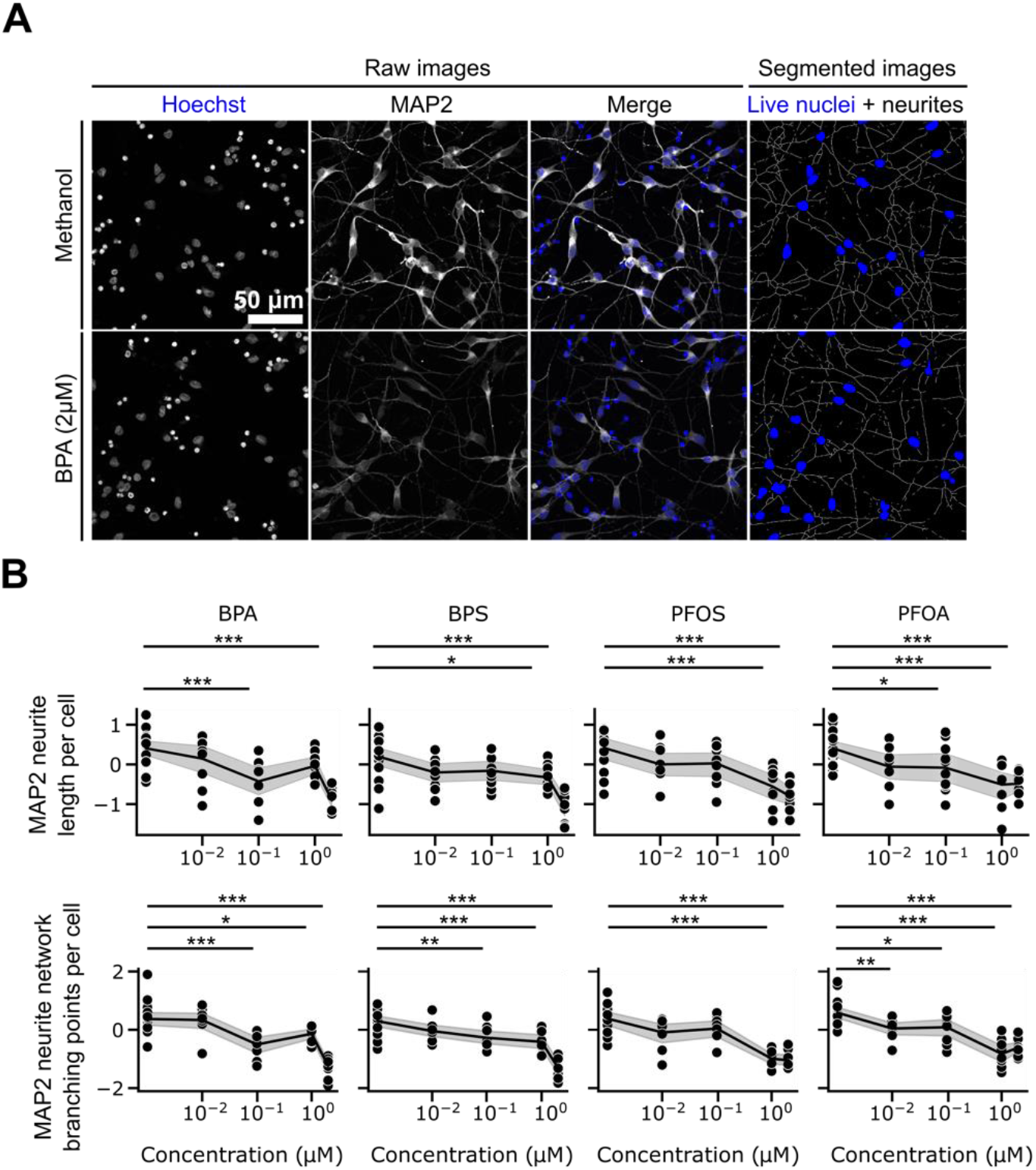
Neurite length and branching points of mDANs are negatively affected by ED treatment. (A) Representative images of methanol control and 2 μM BPA treated mDANs counterstained with Hoechst and a MAP2 antibody. (B) Dose-response graphs and statistical analysis describing the effect of treatments on neurite length per cell, and neurite branching points per cell. Data are shown as normalized single values (black dots), mean (black lines) and 95% CI of the mean (gray area). *p<.05, **p<.01, ***p<.001 compared to methanol control.

In addition to reduced neurite length, the treatments also resulted in a significant decrease in the number of branching points, that are critical for the formation of complex neuronal networks and communication (**Figure 5B**, bottom row). Branching points per cell were significantly decreased when mDANs were exposed to BPA and BPS at 0.1, 1 and 2 μM. PFOS decreased this feature only at 1 and 2 μM, while PFOA had the strongest effect, showing significant differences from 0.01 to 2 μM compared to the methanol control.

### BP and PF exposure increases TH signal intensity and the cellular surface intensity of the TH/αSyn double positive cells

Both classification algorithms showed that ED-treated and untreated mDANs differ also in the membrane-associated TH and αSyn signal mean intensities (**Figure 4D** and **G**). TH and αSyn, whose signal was affected by the xenobiotic treatment (**Figure 6A)** are two crucial proteins in mDANs. A significant fluorescence intensity increase for TH, the enzyme critical for dopamine biosynthesis, was detected inside the neurons: the signal marked the entire cell body, but an increase around the cellular membrane was also observed. Statistical analysis showed that BPA and BPS increased cytoplasmic TH levels at 0.1 and 2 μM. Both PFOS and PFOA lead to increased TH signal intensity in the cytoplasm at 1 and 2 μM, and PFOA also increased this feature at 0.1 μM.

**Figure 6.**
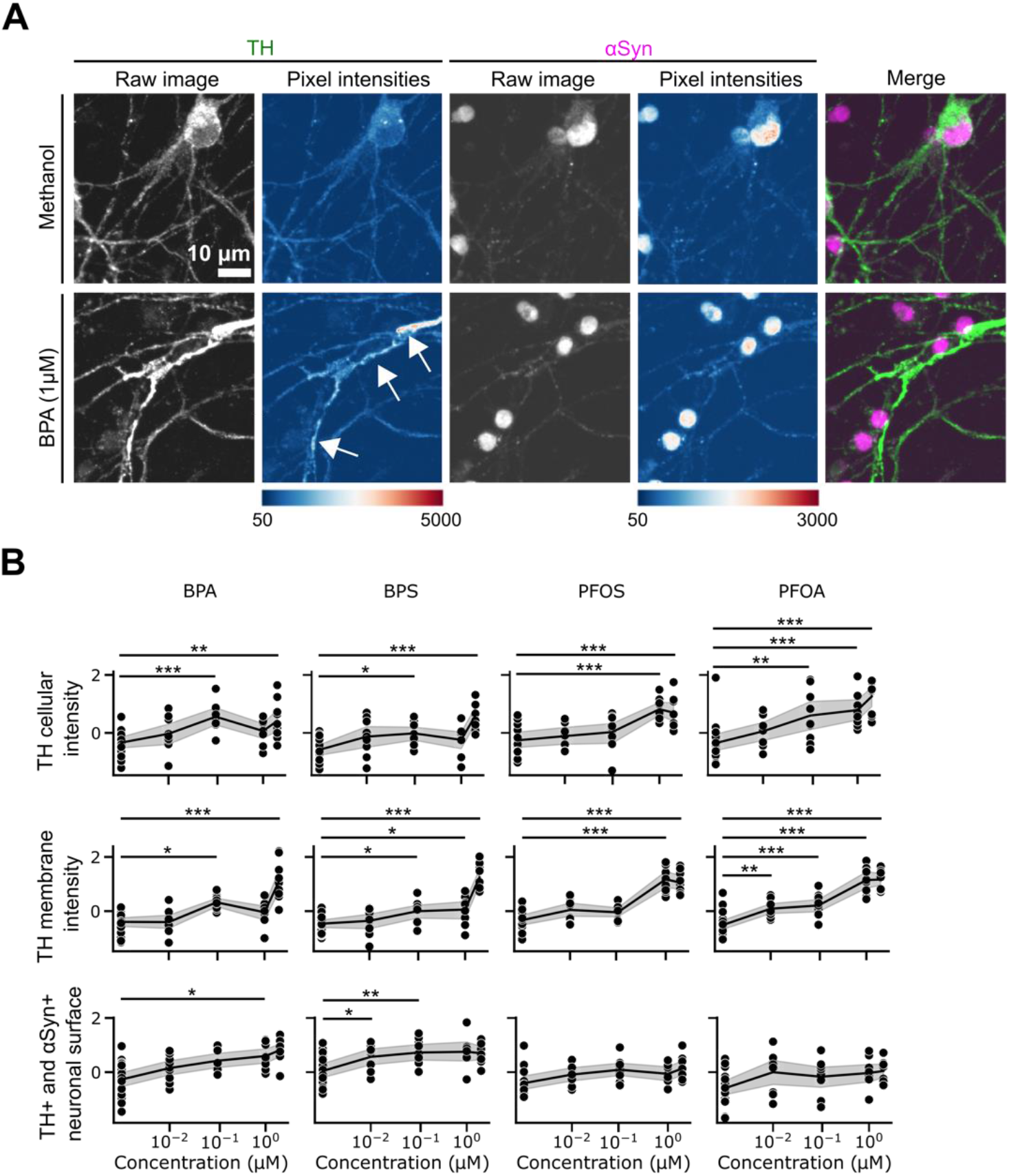
TH signal intensity increases following ED exposure. (A) Representative images of methanol control and 1 μM BPA treated mDANs. Additionally, pixel intensities are illustrated using a heatmap. (B) Dose-response graphs and statistical analysis describing the effect of treatments on overall TH intensity (top panel), around the membrane (middle panel), and surface intensity increase of TH/αSyn double positive cells. Data are shown as normalized single values (black dots), mean (black lines) and 95% CI of the mean (gray area). *p<.05, **p<.01, ***p<.001 compared to methanol control.

Also, BPA significantly increased TH intensity around the membrane at 0.1 and 2 μM. Similarly, BPS increased the TH signal intensity around the membrane at 0.1, 1, and 2 μM. PFOS increased TH intensity around the membrane only at 1 and 2 μM, while PFOA induced this increase at all concentrations compared to control (**Figure 6B**, top and middle panels). We also observed an increase of the cellular surface intensity of TH/αSyn double positive neurons which was elicited only by BPs, and not by PFs. BPA increased the cellular surface intensity of TH/αSyn at 1 μM, while BPS had this effect at 0.01 and 0.1 μM (**Figure 6B**, bottom panel). The increase of TH/αSyn double positive total cellular fluorescence surface intensity is in accordance with the whole cell TH and αSyn intensity increases we described earlier (**Figure 2D**).

Together our findings on MAP2 neurite length and branching points as well as TH and αSyn level changes illustrate how ML classification can aid in the identification of subcellular phenotypes that can be easily overlooked when just visually inspecting images without prior data analysis.

## Discussion

BPs and PFs belong to the ED class due to their ability to interfere with the endocrine system. Recent studies suggest that these compounds can have detrimental effects on the nervous system and that they can worsen PD symptoms in different PD model systems [16– 21], while the effect on human mDANs is unclear. Here, we investigated the effect of these chemicals on human mDANs evaluating their action on protein and morphological features generally affected during PD. The main findings of this study are that i) BPs and PFs lead to a net increase of αSyn protein level, a characteristic hallmark of PD; ii) ED treatment dramatically impaired the neuron network, decreasing neurite length and the branching points per cell; iii) ML successfully classified cells treated with the selected compounds compared to methanol only controls allowing to extract phenotypic features and feature combinations that can be easily overlooked when inspecting cells visually.

Although association studies already suggested a relationship between the exposure to EDs and PD development, our results provide the first evidence of the threatening action of BPs and PFs on a human mDAN model, showing the neuronal biological features affected by EDs exposure. In line with previous data obtained in other neuronal or stem cell models [30– 32], in our setting BPs and PFs did not impact cell viability. However, high content imaging analysis showed that 72-hour exposure to BPA, BPS, or PFOA determined an increase of αSyn levels in human mDANs and this effect was particularly evident at the highest concentration tested (2 μM) (**Figure 2D, Figure 3B, Figure 6B**). Accordingly, Pradyumna and colleagues showed that BPA treatment induced a significant upregulation of both PD-associated αSyn and leucine-rich repeated kinase 2 (LRRK2) proteins in zebrafish [33]. αSyn accumulation in neurons after BPA exposure has also been reported in mammals, being upregulated in the *substantia nigra pars compacta* of adult rats that were neonatally treated [34]. Moreover, when mice were exposed *in utero* to human-relevant doses of BPA, αSyn was one of the most upregulated genes as shown by Ingenuity Pathway Analysis [35]. Effects of BPS and PFs on αSyn levels have not been reported; however, a single oral dose of either PFOS or PFOA increased the levels of tau protein in the cerebral cortex and hippocampus of mice, indicating a role in the dysregulation of normal neural homeostasis. The marked increase in αSyn fluorescence intensity within the mDANs cytoplasm following BPA and BPS exposure is noteworthy, as αSyn plays a central role in the pathogenesis of PD and other synucleinopathies [36]. The elevation of αSyn levels within the cytoplasm suggests altered protein aggregation dynamics or impaired protein degradation mechanisms. These findings are in line with the growing evidence linking environmental exposure to αSyn pathology and neurodegenerative processes [37].

Like for αSyn, alterations of TH fluorescent intensities in specific cell compartments were also identified by both ML classifiers upon the ED exposure (**Figure 4D** and **G, Figure 6**). TH is a critical enzyme involved in dopamine biosynthesis [38]; the TH level increase in both cytoplasm and the neuronal membrane could account for an increased synthesis or for a decreased degradation the enzyme. While the increased signal at the membrane could also suggest an intracellular redistribution from the cytoplasm to the membrane, the overall increased of TH signal after BP treatment seems to rule out this hypothesis, supporting the potential BP effect on TH metabolism (**Figure 6B**, top and middle panels). This finding is consistent with previous studies demonstrating that BPA and BPS may affect dopaminergic pathways and neurotransmitter function [39]. It is known that cellular TH levels decrease during PD, affecting dopamine synthesis [40]. In our setting, the increased levels of TH could reflect an initial compensatory mechanism exerted by mDANs in response to ED exposure. However, the hypothesis that TH signal increase might reflect a higher concentration due to the reduced cellular area determined by the modification of neurite morphology cannot be excluded. We also noted that the overall cellular surface labelled by both TH and αSyn antibodies (TH/αSyn double positive cellular surface intensity) increased following BPA and BPS, but not PFOA or PFOS exposure. These results confirm the neurotoxic effects of BP exposure on human mDANs thus corroborating current literature supporting the role of this ED class in the development of neurodegenerative diseases (**Figure 6B**, bottom panel).

Closer analysis of neurite-related features demonstrated that exposure to BPA, BPS, PFOS, and PFOA negatively impacts neurite length and the number of branching points in mDANs. These findings suggest that these EDs may disrupt the structural development of neuronal processes, which are critical for proper connectivity and communication within neuronal networks [41]. The observed effects exerted by “real life” exposure doses highlight the sensitivity of mDANs to these environmental chemicals, with higher concentrations leading to more significant alterations in neuronal morphology (**Figure 5B**). The BPs and PFs detrimental effects on neurite length and branching points might have implications on neurodevelopmental processes, the proper functioning of dopaminergic circuits and during PD development [42,43]. Moreover, our findings align with previous research demonstrating the neurotoxic effects of BPs on neuronal morphology and connectivity. Indeed, BPA and BPS decreased both normalized neurite total length and normalized maximum neurite length in neuron-like cells at 1 nM while other analogs required higher concentration to exert such a negative effect [44]. Similarly, PFOS and PFOA have been associated with adverse effects on neuronal development and connectivity in other experimental models. Liao *et al*. investigated the effect of different PFs on cultured rat hippocampal neurons and demonstrated that PFOS and PFOA decreased neurite length by about 25% and 20%, respectively [45]. It is also known that BPs and PFs, interfering with the endocrine system, do not show a linear dose-response effect [31,32]. Accordingly, in our study we found that some investigated features were altered at low (0.1 μM) and high concentration (2 μM), but not at medium ones (1 μM).

Taking not only single but all phenotypic features into account, Cosine distance-based clustering and data embedding using UMAP and PaCMAP showed that at 2 μM BPA and BPS phenotypic profiles are similar to each other but differ from PFOA and PFOS profiles which are also similar to each other (**Figure 3B** and **C**). This particular clustering seems to suggest that xenobiotics belonging to the same class of compounds exert similar effects, probably due to the structural similarity and to the consequent ability to interfere with cellular biology in a similar manner.

One of the notable findings of this study is the ability of ML to accurately discriminate and classify the phenotypic profiles of mDANs treated with EDs. Specifically, when the LightGBM algorithm was applied to our image-derived dataset, it correctly classified mDANs treated with EDs and control cells with a high accuracy of 0.92. (**Figure 4F**). We previously reported the effectiveness of ML in classifying normal and PD affected mDANs by applying a Linear Discriminant Analysis (LDA) and Support Vector Machine (SVM) classification to image-derived datasets [25]. ML can help to identify subtle phenotypic patterns that may be easily overlooked but that may be relevant representing the initial signs of cellular distress. This approach appears to be particularly important in the context of the studies on environmental chemicals’ effects on human health. Humans are chronically in contact with these biologically active substances and the consequences on the neurotransmitter systems can become clearly evident only after a protracted exposure, making their risk assessment very challenging. By analyzing large datasets, ML algorithms can detect complex relationships and patterns that may not be immediately apparent to human observers. In this study, by means of the ML approach ED-treated and untreated mDANs were differently classified on the basis of relevant biological features related to PD, such as increased αSyn expression and diminished neurite network length. These data are particularly important, as we exposed human mDANs to BPs and PFs doses that resemble the real-life exposure range.

## Conclusion

Our results provide important information regarding the effect of BPA, BPS, PFOS and PFOA on human mDANs, showing that they drive mDANs toward PD-like phenotypes. Importantly, our ML-supported image analysis approach can identify phenotypic changes that define detrimental EDs effects, thus representing a useful tool for further mechanistic neurotoxicity studies.

## Authors’ contributions

Conception and design: ADC, AW, JHW, ADB; Analysis and interpretation of data: ADC, JHW, MC, GG, BG; Drafting of the manuscript: ADC, JHW; Critical manuscript revision for important intellectual content: ADC, JHW, ADB All co-authors approved the final version of the manuscript.

## References

1. Poewe W, Seppi K, Tanner CM, Halliday GM, Brundin P, Volkmann J, Schrag AE, Lang AE. Parkinson disease. Nat Rev Dis Primers. 2017 Mar 23;3(1):17013.

2. Crowther RA, Daniel SE, Goedert M. Characterisation of isolated alpha-synuclein filaments from substantia nigra of Parkinson’s disease brain. Neurosci Lett. 2000 Oct 6;292(2):128–30.

3. Zhu Y, Zhang J, Zeng Y. Overview of tyrosine hydroxylase in Parkinson’s disease. CNS Neurol Disord Drug Targets. 2012 Jun 1;11(4):350–8.

4. Sveinbjornsdottir S. The clinical symptoms of Parkinson’s disease. J Neurochem. 2016 Oct;139 Suppl 1:318–24.

5. Kumar M, Sarma DK, Shubham S, Kumawat M, Verma V, Prakash A, Tiwari R. Environmental Endocrine-Disrupting Chemical Exposure: Role in Non-Communicable Diseases. Front Public Health. 2020 Sep 24;8:553850.

6. Ješeta M, Navrátilová J, Franzová K, Fialková S, Kempisty B, Ventruba P, žáková J, Crha I. Overview of the Mechanisms of Action of Selected Bisphenols and Perfluoroalkyl Chemicals on the Male Reproductive Axes. Front Genet. 2021 Sep 27;12:692897.

7. D’Hollander W, de Voogt P, De Coen W, Bervoets L. Perfluorinated substances in human food and other sources of human exposure. Rev Environ Contam Toxicol. 2010;208:179–215.

8. Geens T, Aerts D, Berthot C, Bourguignon JP, Goeyens L, Lecomte P, Maghuin-Rogister G, Pironnet AM, Pussemier L, Scippo ML, Van Loco J, Covaci A. A review of dietary and non-dietary exposure to bisphenol-A. Food Chem Toxicol. 2012 Oct;50(10):3725–40.

9. Jin H, Xie J, Mao L, Zhao M, Bai X, Wen J, Shen T, Wu P. Bisphenol analogue concentrations in human breast milk and their associations with postnatal infant growth. Environmental Pollution. 2020 Apr;259:113779.

10. Lee J, Choi K, Park J, Moon HB, Choi G, Lee JJ, Suh E, Kim HJ, Eun SH, Kim GH, Cho GJ, Kim SK, Kim S, Kim SY, Kim S, Eom S, Choi S, Kim YD, Kim S. Bisphenol A distribution in serum, urine, placenta, breast milk, and umbilical cord serum in a birth panel of mother–neonate pairs. Science of The Total Environment. 2018 Jun;626:1494–501.

11. Hall SM, Zhang S, Hoffman K, Miranda ML, Stapleton HM. Concentrations of per- and polyfluoroalkyl substances (PFAS) in human placental tissues and associations with birth outcomes. Chemosphere. 2022 May;295:133873.

12. Teng C, Goodwin B, Shockley K, Xia M, Huang R, Norris J, Merrick BA, Jetten AM, Austin CP, Tice RR. Bisphenol A affects androgen receptor function via multiple mechanisms. Chemico-Biological Interactions. 2013 May;203(3):556–64.

13. Du G, Hu J, Huang H, Qin Y, Han X, Wu D, Song L, Xia Y, Wang X. Perfluorooctane sulfonate (PFOS) affects hormone receptor activity, steroidogenesis, and expression of endocrine-related genes in vitro and in vivo. Environmental Toxicology and Chemistry. 2013 Feb;32(2):353–60.

14. Vandenberg LN, Colborn T, Hayes TB, Heindel JJ, Jacobs DR, Lee DH, Shioda T, Soto AM, Vom Saal FS, Welshons WV, Zoeller RT, Myers JP. Hormones and Endocrine-Disrupting Chemicals: Low-Dose Effects and Nonmonotonic Dose Responses. Endocrine Reviews. 2012 Jun 1;33(3):378–455.

15. Schug TT, Janesick A, Blumberg B, Heindel JJ. Endocrine disrupting chemicals and disease susceptibility. The Journal of Steroid Biochemistry and Molecular Biology. 2011 Nov;127(3–5):204–15.

16. Gu J, Guo M, Yin X, Huang C, Qian L, Zhou L, Wang Z, Wang L, Shi L, Ji G. A systematic comparison of neurotoxicity of bisphenol A and its derivatives in zebrafish. Science of The Total Environment. 2022 Jan;805:150210.

17. Tukker AM, Bouwman LMS, van Kleef RGDM, Hendriks HS, Legler J, Westerink RHS. Perfluorooctane sulfonate (PFOS) and perfluorooctanoate (PFOA) acutely affect human α1β2γ2L GABAA receptor and spontaneous neuronal network function in vitro. Sci Rep. 2020 Mar 24;10(1):5311.

18. Inadera H. Neurological Effects of Bisphenol A and its Analogues. Int J Med Sci. 2015;12(12):926–36.

19. Musachio EAS, Araujo SM, Bortolotto VC, de Freitas Couto S, Dahleh MMM, Poetini MR, Jardim EF, Meichtry LB, Ramborger BP, Roehrs R, Petri Guerra G, Prigol M. Bisphenol A exposure is involved in the development of Parkinson like disease in Drosophila melanogaster. Food and Chemical Toxicology. 2020 Mar;137:111128.

20. Elsworth JD, Jentsch JD, VandeVoort CA, Roth RH, Jr DER Leranth C. Prenatal exposure to bisphenol A impacts midbrain dopamine neurons and hippocampal spine synapses in non-human primates. NeuroToxicology. 2013 Mar;35:113–20.

21. D’Amico R, Gugliandolo E, Siracusa R, Cordaro M, Genovese T, Peritore AF, Crupi R, Interdonato L, Di Paola D, Cuzzocrea S, Fusco R, Impellizzeri D, Di Paola R. Toxic Exposure to Endocrine Disruptors Worsens Parkinson’s Disease Progression through NRF2/HO-1 Alteration. Biomedicines. 2022 May 5;10(5):1073.

22. Landolfi A, Troisi J, Savanelli MC, Vitale C, Barone P, Amboni M. Bisphenol A glucuronidation in patients with Parkinson’s disease. NeuroToxicology. 2017 Dec;63:90–6.

23. Qian Y, Ducatman A, Ward R, Leonard S, Bukowski V, Lan Guo N, Shi X, Vallyathan V, Castranova V. Perfluorooctane Sulfonate (PFOS) Induces Reactive Oxygen Species (ROS) Production in Human Microvascular Endothelial Cells: Role in Endothelial Permeability. Journal of Toxicology and Environmental Health, Part A. 2010 Apr 14;73(12):819–36.

24. Nakagawa Y, Tayama S. Metabolism and cytotoxicity of bisphenol A and other bisphenols in isolated rat hepatocytes. Archives of Toxicology. 2000 Apr 25;74(2):99–105.

25. Vuidel A, Cousin L, Weykopf B, Haupt S, Hanifehlou Z, Wiest-Daesslé N, Segschneider M, Lee J, Kwon YJ, Peitz M, Ogier A, Brino L, Brüstle O, Sommer P, Wilbertz JH. High-content phenotyping of Parkinson’s disease patient stem cell-derived midbrain dopaminergic neurons using machine learning classification. Stem Cell Reports. 2022 Oct;17(10):2349–64.

26. Monzel AS, Hemmer K, Kaoma T, Smits LM, Bolognin S, Lucarelli P, Rosety I, Zagare A, Antony P, Nickels SL, Krueger R, Azuaje F, Schwamborn JC. Machine learning-assisted neurotoxicity prediction in human midbrain organoids. Parkinsonism & Related Disorders. 2020 Jun;75:105–9.

27. Schwartz MP, Hou Z, Propson NE, Zhang J, Engstrom CJ, Costa VS, Jiang P, Nguyen BK, Bolin JM, Daly W, Wang Y, Stewart R, Page CD, Murphy WL, Thomson JA. Human pluripotent stem cell-derived neural constructs for predicting neural toxicity. Proc Natl Acad Sci USA. 2015 Oct 6;112(40):12516–21.

28. McInnes L, Healy J, Melville J. UMAP: Uniform Manifold Approximation and Projection for Dimension Reduction. 2018 [cited 2023 Jul 31]; Available from: https://arxiv.org/abs/1802.03426

29. Wang Y, Huang H, Rudin C, Shaposhnik Y. Understanding How Dimension Reduction Tools Work: An Empirical Approach to Deciphering t-SNE, UMAP, TriMAP, and PaCMAP for Data Visualization. 2020 [cited 2023 Jul 31]; Available from: https://arxiv.org/abs/2012.04456

30. Pierozan P, Karlsson O. Differential susceptibility of rat primary neurons and neural stem cells to PFOS and PFOA toxicity. Toxicology Letters. 2021 Oct;349:61–8.

31. Gaggi G, Di Credico A, Barbagallo F, Ballerini P, Ghinassi B, Di Baldassarre A. Antenatal Exposure to Plastic Pollutants: Study of the Bisphenols and Perfluoroalkyls Effects on Human Stem Cell Models. Expo Health [Internet]. 2023 Jul 18 [cited 2023 Aug 8]; Available from: https://link.springer.com/10.1007/s12403-023-00586-5

32. Gaggi G, Di Credico A, Barbagallo F, Ghinassi B, Di Baldassarre A. Bisphenols and perfluoroalkyls alter human stem cells integrity: A possible link with infertility. Environmental Research. 2023 Oct;235:116487.

33. Sahoo PK, Aparna S, Naik PK, Singh SB, Das SK. Bisphenol A exposure induces neurobehavioral deficits and neurodegeneration through induction of oxidative stress and activated caspase‐3 expression in zebrafish brain. J Biochem Mol Toxicol [Internet]. 2021 Oct [cited 2023 Jul 31];35(10). Available from: https://onlinelibrary.wiley.com/doi/10.1002/jbt.22873

34. Ishido M, Masuo Y. Temporal Effects of Bisphenol A on Dopaminergic Neurons: An Experiment on Adult Rats. TOENVIRJ. 2014 May 16;8(1):9–17.

35. Henriksen AD, Andrade A, Harris EP, Rissman EF, Wolstenholme JT. Bisphenol A Exposure in utero Disrupts Hypothalamic Gene Expression Particularly Genes Suspected in Autism Spectrum Disorders and Neuron and Hormone Signaling. IJMS. 2020 Apr 29;21(9):3129.

36. Calabresi P, Mechelli A, Natale G, Volpicelli-Daley L, Di Lazzaro G, Ghiglieri V. Alpha-synuclein in Parkinson’s disease and other synucleinopathies: from overt neurodegeneration back to early synaptic dysfunction. Cell Death Dis. 2023 Mar 1;14(3):176.

37. Rebolledo-Solleiro D. Impact of BPA on behavior, neurodevelopment and neurodegeneration Daniela. Front Biosci. 2021;26(2):363–400.

38. Meiser J, Weindl D, Hiller K. Complexity of dopamine metabolism. Cell Commun Signal. 2013 Dec;11(1):34.

39. Wang Y, Gai T, Zhang L, Chen L, Wang S, Ye T, Zhang W. Neurotoxicity of bisphenol A exposure on Caenorhabditis elegans induced by disturbance of neurotransmitter and oxidative damage. Ecotoxicology and Environmental Safety. 2023 Mar;252:114617.

40. Cramb KML, Beccano-Kelly D, Cragg SJ, Wade-Martins R. Impaired dopamine release in Parkinson’s disease. Brain. 2023 Mar 2;awad064.

41. Kulkarni VA, Firestein BL. The dendritic tree and brain disorders. Molecular and Cellular Neuroscience. 2012 May;50(1):10–20.

42. Mainen ZF, Sejnowski TJ. Influence of dendritic structure on firing pattern in model neocortical neurons. Nature. 1996 Jul;382(6589):363–6.

43. Maiti P, Manna J, Ilavazhagan G, Rossignol J, Dunbar GL. Molecular regulation of dendritic spine dynamics and their potential impact on synaptic plasticity and neurological diseases. Neuroscience & Biobehavioral Reviews. 2015 Dec;59:208–37.

44. Liang X, Yin N, Liang S, Yang R, Liu S, Lu Y, Jiang L, Zhou Q, Jiang G, Faiola F. Bisphenol A and several derivatives exert neural toxicity in human neuron-like cells by decreasing neurite length. Food and Chemical Toxicology. 2020 Jan;135:111015.

45. Liao C, Wang T, Cui L, Zhou Q, Duan S, Jiang G. Changes in Synaptic Transmission, Calcium Current, and Neurite Growth by Perfluorinated Compounds Are Dependent on the Chain Length and Functional Group. Environ Sci Technol. 2009 Mar 15;43(6):2099–104.

