## Supplemental Table 1 for "Machine learning identifies phenotypic profile alterations of human dopaminergic neurons exposed to bisphenols and perfluoroalkyls"

| Name of Material/ Equipment | Company | Catalog Number |
|---|---|---|
| Anti- chicken – Alexa 647 | Jackson ImmunoRearch | 703-605-155 |
| Anti-Map2 | Novus | NB300-213 |
| Anti-mouse - Alexa 488 | Thermo Fisher | A11001 |
| Anti-rabbit - Alexa 555 | Thermo Fisher | A21429 |
| Anti-Tyrosine Hydroxylase | Merck | T2928 |
| Anti-α-synuclein | Abcam | 138501 |
| Bisphenol A (BPA) | Wellington Laboratories | BPA (80-05-7) |
| Bisphenol S (BPS) | Wellington Laboratories | BPS (80-09-1) |
| Bravo Automated Liquid Handling Platform with 384ST head | Agilent | |
| Confocal microscope | Yokogawa | CV7000 |
| Countess Automated cell counter | Invitrogen | |
| DPBS +/+ | Gibco | 14040-133 |
| EL406 Washer Dispenser | BioTek (Agilent) | |
| Formaldehyde Solution (PFA 16 %) | Euromedex | EM-15710-S |
| Hoechst 33342 | Invitrogen | H3570 |
| iCell Base Medium 1 | Fujifilm | M1010 |
| iCell DPN, Donor#01279, Phenotype AHN, lot#106339, 1M | Fujifilm | C1087 |
| iCell Nervous System Supplement | Fujifilm | M1031 |
| iCell Neural Supplement B | Fujifilm | M1029 |
| Jupyter Python notebook for data analysis and plotting | In-house development | https://github.com/Ksilink/Notebooks/tree/main/Neuro/EndocrineDisruptorProfiling |
| Laminin | Biolamina | LN521 |
| Perfluorooctane sulfonate (PFOS) | Wellington Laboratories | L-PFOS (4021-47-0) |
| Perfluorooctanoic acid (PFOA) | Wellington Laboratories | PFOA (335-67-1) |
| PFE-360 | MedChemExpress | HY-120085 |
| PhenoLink image segmentation software | In-house development | https://github.com/Ksilink/PhenoLink |
| PhenoPlate 384w, PDL coated | Perkin Elmer | 6057500 |
| Storage plates Abgene 120µL | Thermo Scientific | AB-0781 |
| Triton | Sigma | T9284 |
| Trypan Blue | Sigma | T8154-20ML |
| Vprep Pipetting System | Agilent | |
