## Supplemental Table 2 for "Machine learning identifies phenotypic profile alterations of human dopaminergic neurons exposed to bisphenols and perfluoroalkyls"

| Complete Maintenance Media | |
|---|---|
| **Reagents** | **Final conc.** |
| iCell Base Medium 1 | 0.97 |
| iCell Neural Supplement B | 0.02 |
| iCell Nervous System Suppl. | 0.01 |
| **Blocking solution** | |
| **Reagent** | **Final conc.** |
| PBS 1X | 1 |
| Triton 10 % | 0.001 |
| FBS stock solution | 0.1 |
| **Primary staining solution** | |
| **Reagent** | **Final conc.** |
| PBS 1X | 1 |
| Triton 10 % | 0.001 |
| FBS stock solution | 0.05 |
| Anti-TH | 1/1000 |
| Anti-α-synuclein | 1/500 |
| Anti-MAP2 | 1/5000 |
| **Secondary staining solution** | |
| **Reagent** | **Final conc.** |
| PBS 1X | 1 |
| Triton 10 % | 0.001 |
| FBS stock solution | 0.05 |
| Anti-mouse A488 | 1/1000 |
| Anti-rabbit A555 | 1/1000 |
| Anti-chicken A647 | 1/250 |
| Hoechst | 1/2000 |
