## Supplemental Table 3 for "Machine learning identifies phenotypic profile alterations of human dopaminergic neurons exposed to bisphenols and perfluoroalkyls"

| | Feature name | Feature description |
|---|---|---|
| 1 | Cell_Correlation_MAP2_SNCA | Pearson correlation between MAP2 and SNCA channel |
| 2 | Cell_Correlation_TH_SNCA | Pearson correlation between TH and SNCA channel |
| 3 | Cell_Correlation_VanSteelselsMeanX_TH_SNCA | Van Steelsel's cross correlation between TH and SNCA channel, shift on x-axis |
| 4 | Cell_Correlation_VanSteelselsMeanY_TH_SNCA | Van Steelsel's cross correlation between TH and SNCA channel, shift on y-axis |
| 5 | Cell_Correlation_VanSteelselsSigmaX_TH_SNCA | SD of Van Steelsel's cross correlation between TH and SNCA channel, shift on x-axis |
| 6 | Cell_Correlation_VanSteelselsSigmaY_TH_SNCA | SD of Van Steelsel's cross correlation between TH and SNCA channel, shift on y-axis |
| 7 | Cell_Intensity_MeanIntensity_MAP2 | Mean pixel intensity of MAP2 channel |
| 8 | Cell_Intensity_MeanIntensity_SNCA | Mean pixel intensity of SNCA channel |
| 9 | Cell_Intensity_MeanIntensity_TH | Mean pixel intensity of TH channel |
| 10 | Cell_Intensity_SumIntensityPerNuclei_SNCA | Integrated pixel intensity of SNCA channel normalized to number of nuclei |
| 11 | Cell_MAP2_SNCA_Intensity_MeanIntensity_SNCA | Mean pixel intensity of SNCA channel colocalized to MAP2 channel |
| 12 | Cell_Neurites_BranchingPointsPerNuclei_MAP2 | Dendritic branching points of MAP2 channel normalized to number of nuclei |
| 13 | Cell_Neurites_BranchingPointsPerNuclei_TH | Dendritic branching points of TH channel normalized to number of nuclei |
| 14 | Cell_Neurites_LengthPerNuclei_MAP2 | Dentritic network length of MAP2 channel normalized to number of nuclei |
| 15 | Cell_Neurites_LengthPerNuclei_TH | Dentritic network length of TH channel normalized to number of nuclei |
| 16 | Cell_Neurites_Length_MAP2 | Dentritic network length of MAP2 channel |
| 17 | Cell_Neurites_Length_TH | Dentritic network length of TH channel |
| 18 | Cell_SurfacePerNuclei_MAP2 | Surface pixels occupied by MAP2 channel, normalized to number of nuclei |
| 19 | Cell_SurfacePerNuclei_MAP2_SNCA | Surface pixels occupied by colocalized MAP2 and SNCA channel normalized to number of nuclei |
| 20 | Cell_SurfacePerNuclei_SNCA | Surface pixels occupied by SNCA channel, normalized to number of nuclei |
| 21 | Cell_SurfacePerNuclei_TH | Surface pixels occupied by TH channel, normalized to number of nuclei |
| 22 | Cell_SurfacePerNuclei_TH_SNCA | Surface pixels occupied by colocalized TH and SNCA channel normalized to number of nuclei |
| 23 | Cell_Surface_RatioSurface_TH_SNCA | Surface ratio occupied by colocalized TH and SNCA channel |
| 24 | Cell_Surface_TotalSurface_MAP2 | Surface pixels occupied by MAP2 channel |
| 25 | Cell_Surface_TotalSurface_SNCA | Surface pixels occupied by SNCA channel |
| 26 | Cell_Surface_TotalSurface_SNCA_MAP2 | Surface pixels occupied by colocalized SNCA and MAP2 channel |
| 27 | Cell_Surface_TotalSurface_TH | Surface pixels occupied by TH channel |
| 28 | Cell_Surface_TotalSurface_TH_SNCA | Surface pixels occupied by colocalized TH and SNCA channel |
| 29 | Cell_TH_SNCA_Intensity_MeanIntensity_SNCA | Mean pixel intensity of SNCA channel colocalized to TH channel |
| 30 | Cell_TH_SNCA_Intensity_MeanIntensity_TH | Mean pixel intensity of TH channel colocalized to SNCA channel |
| 31 | Cell_TH_SNCA_Intensity_SumIntensityPerNuclei_SNCA | Integrated pixel intensity of SNCA channel colocalized to TH channel normalized to number of nuclei |
| 32 | Cell_Texture_SNCA_AngularSecondMoment_000 | Haralick uniformity of distribution of gray levels at 0 degree shift |
| 33 | Cell_Texture_SNCA_AngularSecondMoment_045 | Haralick uniformity of distribution of gray levels at 45 degree shift |
| 34 | Cell_Texture_SNCA_AngularSecondMoment_090 | Haralick uniformity of distribution of gray levels at 90 degree shift |
| 35 | Cell_Texture_SNCA_AngularSecondMoment_135 | Haralick uniformity of distribution of gray levels at 135 degree shift |
| 36 | Cell_Texture_SNCA_Contrast_000 | Haralick contrast of gray levels at 0 degree shift |
| 37 | Cell_Texture_SNCA_Contrast_045 | Haralick contrast of gray levels at 45 degree shift |
| 38 | Cell_Texture_SNCA_Contrast_090 | Haralick contrast of gray levels at 90 degree shift |
| 39 | Cell_Texture_SNCA_Contrast_135 | Haralick contrast of gray levels at 135 degree shift |
| 40 | Cell_Texture_SNCA_Correlation_000 | Haralick correlation of gray levels at 0 degree shift |
| 41 | Cell_Texture_SNCA_Correlation_045 | Haralick correlation of gray levels at 45 degree shift |
| 42 | Cell_Texture_SNCA_Correlation_090 | Haralick correlation of gray levels at 90 degree shift |
| 43 | Cell_Texture_SNCA_Correlation_135 | Haralick correlation of gray levels at 135 degree shift |
| 44 | Cell_Texture_SNCA_DifferenceEntropy_000 | Haralick difference of randomness of gray levels at 0 degree shift |
| 45 | Cell_Texture_SNCA_DifferenceEntropy_045 | Haralick difference of randomness of gray levels at 45 degree shift |
| 46 | Cell_Texture_SNCA_DifferenceEntropy_090 | Haralick difference of randomness of gray levels at 90 degree shift |
| 47 | Cell_Texture_SNCA_DifferenceEntropy_135 | Haralick difference of randomness of gray levels at 135 degree shift |
| 48 | Cell_Texture_SNCA_DifferenceVariance_000 | Haralick difference of variance of gray levels at 0 degree shift |
| 49 | Cell_Texture_SNCA_DifferenceVariance_045 | Haralick difference of variance randomness of gray levels at 45 degree shift |
| 50 | Cell_Texture_SNCA_DifferenceVariance_090 | Haralick difference of variance randomness of gray levels at 90 degree shift |
| 51 | Cell_Texture_SNCA_DifferenceVariance_135 | Haralick difference of variance randomness of gray levels at 135 degree shift |
| 52 | Cell_Texture_SNCA_Entropy_000 | Haralick randomness of gray levels at 0 degree shift |
| 53 | Cell_Texture_SNCA_Entropy_045 | Haralick randomness of gray levels at 45 degree shift |
| 54 | Cell_Texture_SNCA_Entropy_090 | Haralick randomness of gray levels at 90 degree shift |
| 55 | Cell_Texture_SNCA_Entropy_135 | Haralick randomness of gray levels at 135 degree shift |
| 56 | Cell_Texture_SNCA_InfoMeasuresOfCorr1_000 | Haralick information measure of correlation 1 of gray levels at 0 degree shift |
| 57 | Cell_Texture_SNCA_InfoMeasuresOfCorr1_045 | Haralick information measure of correlation 1 of gray levels at 45 degree shift |
| 58 | Cell_Texture_SNCA_InfoMeasuresOfCorr1_090 | Haralick information measure of correlation 1 of gray levels at 90 degree shift |
| 59 | Cell_Texture_SNCA_InfoMeasuresOfCorr1_135 | Haralick information measure of correlation 1 of gray levels at 135 degree shift |
| 60 | Cell_Texture_SNCA_InfoMeasuresOfCorr2_000 | Haralick information measure of correlation 2 of gray levels at 0 degree shift |
| 61 | Cell_Texture_SNCA_InfoMeasuresOfCorr2_045 | Haralick information measure of correlation 2 of gray levels at 45 degree shift |
| 62 | Cell_Texture_SNCA_InfoMeasuresOfCorr2_090 | Haralick information measure of correlation 2 of gray levels at 90 degree shift |
| 63 | Cell_Texture_SNCA_InfoMeasuresOfCorr2_135 | Haralick information measure of correlation 2 of gray levels at 135 degree shift |
| 64 | Cell_Texture_SNCA_InverseDiffMoment_000 | Haralick homogeneity of gray levels at 0 degree shift |
| 65 | Cell_Texture_SNCA_InverseDiffMoment_045 | Haralick homogeneity of gray levels at 45 degree shift |
| 66 | Cell_Texture_SNCA_InverseDiffMoment_090 | Haralick homogeneity of gray levels at 90 degree shift |
| 67 | Cell_Texture_SNCA_InverseDiffMoment_135 | Haralick homogeneity of gray levels at 135 degree shift |
| 68 | Cell_Texture_SNCA_SumAverage_000 | Haralick sum of averages of gray levels at 0 degree shift |
| 69 | Cell_Texture_SNCA_SumAverage_045 | Haralick sum of averages of gray levels at 45 degree shift |
| 70 | Cell_Texture_SNCA_SumAverage_090 | Haralick sum of averages of gray levels at 90 degree shift |
| 71 | Cell_Texture_SNCA_SumAverage_135 | Haralick sum of averages of gray levels at 135 degree shift |
| 72 | Cell_Texture_SNCA_SumEntropy_000 | Haralick sum of gray level randomness at 0 degree shift |
| 73 | Cell_Texture_SNCA_SumEntropy_045 | Haralick sum of gray level randomness at 45 degree shift |
| 74 | Cell_Texture_SNCA_SumEntropy_090 | Haralick sum of gray level randomness at 90 degree shift |
| 75 | Cell_Texture_SNCA_SumEntropy_135 | Haralick sum of gray level randomness at 135 degree shift |
| 76 | Cell_Texture_SNCA_SumOfSquares_000 | Haralick sum of square gray level variance at 0 degree shift |
| 77 | Cell_Texture_SNCA_SumOfSquares_045 | Haralick sum of square gray level variance at 45 degree shift |
| 78 | Cell_Texture_SNCA_SumOfSquares_090 | Haralick sum of square gray level variance at 90 degree shift |
| 79 | Cell_Texture_SNCA_SumOfSquares_135 | Haralick sum of square gray level variance at 135 degree shift |
| 80 | Cell_Texture_SNCA_SumVariance_000 | Haralick sum of gray level variance at 0 degree shift |
| 81 | Cell_Texture_SNCA_SumVariance_045 | Haralick sum of gray level variance at 45 degree shift |
| 82 | Cell_Texture_SNCA_SumVariance_090 | Haralick sum of gray level variance at 90 degree shift |
| 83 | Cell_Texture_SNCA_SumVariance_135 | Haralick sum of gray level variance at 135 degree shift |
| 84 | Cytoplasm_Intensity_MeanIntensity_SNCA | Mean pixel intensity of cytoplasmic SNCA channel |
| 85 | Cytoplasm_MAP2_SNCA_Intensity_MeanIntensity_SNCA | Mean pixel intensity of cytoplasmic SNCA channel colocalized to MAP2 channel |
| 86 | Cytoplasm_SurfacePerNuclei_SNCA | Surface pixels occupied by cytoplasmic SNCA channel normalized to number of nuclei |
| 87 | Cytoplasm_SurfacePerNuclei_TH_SNCA | Surface pixels occupied by colocalized cytoplasmic TH and SNCA channel normalized to number of nuclei |
| 88 | Cytoplasm_Surface_TotalSurface_SNCA | Surface pixels occupied by cytoplasmic SNCA channel |
| 89 | IndividualCell_Intensity_MeanIntensity_SNCA | Mean pixel intensity of SNCA channel based on all individually segmented cells |
| 90 | IndividualCell_Intensity_RadialProfile_InterceptFit_SNCA | Fitted intercept of SNCA channel decay from center to edge |
| 91 | IndividualCell_Intensity_RadialProfile_MaxSlope_SNCA | Maximum steepness of SNCA channel decay from center to edge |
| 92 | IndividualCell_Intensity_RadialProfile_Maximum_SNCA | Maximum intensity of SNCA channel from center to edge |
| 93 | IndividualCell_Intensity_RadialProfile_MeanCoeffVar_SNCA | Mean SNCA channel dispersion from center to edge |
| 94 | IndividualCell_Intensity_RadialProfile_MeanGradient_SNCA | Mean shape of SNCA channel decay from center to edge |
| 95 | IndividualCell_Intensity_RadialProfile_Mean_SNCA | Mean intensity of SNCA channel from center to edge |
| 96 | IndividualCell_Intensity_RadialProfile_Median_SNCA | Median intensity of SNCA channel from center to edge |
| 97 | IndividualCell_Intensity_RadialProfile_Minimum_SNCA | Minimum intensity of SNCA channel from center to edge |
| 98 | IndividualCell_Intensity_RadialProfile_Q1_SNCA | First quartile intensity of SNCA channel from center to edge |
| 99 | IndividualCell_Intensity_RadialProfile_Q3_SNCA | Third quartile intensity of SNCA channel from center to edge |
| 100 | IndividualCell_Intensity_RadialProfile_SlopeFit_SNCA | Fitted slope of SNCA channel decay from center to edge |
| 101 | IndividualCell_Intensity_RadialProfile_StdCoeffVar_SNCA | SD of SNCA channel dispersion from center to edge |
| 102 | IndividualCell_Intensity_RadialProfile_Std_SNCA | SD of SNCA channel intensity from center to edge |
| 103 | IndividualCell_Intensity_SumIntensity_SNCA | Integrated pixel intensity of SNCA channel based on all individually segmented cells |
| 104 | IndividualCell_Surface_MeanSurface_SNCA | Mean surface pixels occupied by SNCA channel based on all individually segmented cells |
| 105 | Membrane_Intensity_MeanIntensity_MAP2 | Mean pixel intensity of MAP2 channel on cellular edge |
| 106 | Membrane_Intensity_MeanIntensity_SNCA | Mean pixel intensity of TH channel on cellular edge |
| 107 | Membrane_Intensity_MeanIntensity_TH | Mean pixel intensity of MAP2 channel on cellular edge |
| 108 | Membrane_Surface_SurfacePerNuclei_MAP2 | Surface pixels on cellular edge occupied by MAP2 channel normalized to number of nuclei |
| 109 | Membrane_Surface_SurfacePerNuclei_SNCA | Surface pixels on cellular edge occupied by SNCA channel normalized to number of nuclei |
| 110 | Membrane_Surface_SurfacePerNuclei_TH | Surface pixels on cellular edge occupied by TH channel normalized to number of nuclei |
| 111 | Nuclei_Living_Ratio_MAP2 | Ratio of MAP2 channel positive nuclei |
| 112 | Nuclei_Living_Ratio_MAP2_SNCA | Ratio of MAP2 and SNCA channel positive nuclei |
| 113 | Nuclei_Living_Ratio_SNCA | Ratio of SNCA channel positive nuclei |
| 114 | Nuclei_Living_Ratio_TH | Ratio of TH channel positive nuclei |
| 115 | Nuclei_Living_Ratio_TH_SNCA | Ratio of TH and SNCA channel positive nuclei |
| 116 | Nuclei_Number_Big | Number of large nuclei |
| 117 | Nuclei_Number_Dead | Number of condensed/bright nuclei |
| 118 | Nuclei_Number_Living | Number of nuclei based on Hoechst channel |
| 119 | Nuclei_Number_MAP2 | Number of MAP2 channel positive nuclei |
| 120 | Nuclei_Number_MAP2_SNCA | Number of MAP2 and SNCA channel positive nuclei |
| 121 | Nuclei_Number_SNCA | Number of SNCA channel positive nuclei |
| 122 | Nuclei_Number_TH | Number of TH channel positive nuclei |
| 123 | Nuclei_Number_TH_SNCA | Number of TH and SNCA channel positive nuclei |
| 124 | Nuclei_Ratio_Dead | Ratio of condensed/bright nuclei |
| 125 | Nuclei_Ratio_Living | Ratio of nuclei not considered condensed/bright |
| 126 | Nuclei_Surface_MeanArea | Mean surface pixels of Hoechst channel |
